# Accumulation and light-harvesting function of IsiA in cyanobacterial cells with monomeric and trimeric Photosystem I

**DOI:** 10.1101/2023.07.12.548727

**Authors:** Parveen Akhtar, Fanny Balog-Vig, Soujanya Kuntam, Szilvia Zita Tóth, Petar H. Lambrev

## Abstract

The acclimation of cyanobacteria to iron deficiency is crucial for their survival in natural environments. In response to iron deficiency, many cyanobacterial species induce the production of a pigment-protein complex called IsiA. IsiA proteins associate with photosystem I (PSI) and can function as light-harvesting antennas or dissipate excess energy. They may also serve as Chl storage during iron limitation. In this study we examined the functional role of IsiA in cells of *Synechocystis* sp. PCC 6803 grown under iron limitation conditions by measuring the cellular IsiA content and its capability to transfer energy to PSI. We specifically test the effect of the oligomeric state of PSI by comparing wild-type (WT) *Synechocystis* sp. PCC 6803 to mutants lacking specific subunits of PSI, namely PsaL/PsaI (Δ*psaL* mutant) and PsaF/PsaJ (ΔFIJL). Time-resolved fluorescence spectroscopy revealed that IsiA formed functional PSI_3_-IsiA_18_ supercomplexes, wherein IsiA effectively transfers energy to PSI on a timescale of 10 ps at room temperature – measured in isolated complexes and in vivo – confirming the primary role of IsiA as an accessory light-harvesting antenna to PSI. However, a significant fraction (40%) remained unconnected to PSI, supporting the notion of a dual functional role of IsiA. Cells with monomeric PSI under iron deficiency contained only 3–4 IsiA complexes bound to PSI. Together the results show that IsiA is capable of transferring energy to trimeric and monomeric PSI but to varying degrees and that the acclimatory production of IsiA under iron stress is controlled by its ability to perform its light-harvesting function.

## INTRODUCTION

The light reactions of photosynthesis in cyanobacteria are carried out in the thylakoid membranes by the two photosystems and the intersystem electron carriers. PSI and PSII contain light-harvesting antenna pigments in their core antennas; in addition, a major fraction of the light energy is captured by the membrane associated phycobilisomes (PBS) that transfer energy captured by the phycobiliproteins to chlorophylls (Chls) in the core antenna. Both the physical association and the energy transfer efficiency between the light-harvesting antenna and the photosystem core complexes can change in response to changes in the environment (Blankenship, 2021).

Iron is a particularly important element for cyanobacteria – serving as a cofactor in various metalloenzymes, including protein complexes involved in respiration, nitrogen fixation, and especially photosynthesis. The photosynthetic apparatus is particularly iron-rich, requiring around 20 iron atoms in the linear electron transport chain (Jia et al., 2021). Therefore, iron deficiency severely hinders photoinduced electron transfer and photosynthetic activity. As iron deficiency is a common nutrient stress in cyanobacterial habitats (Keren et al., 2004), cyanobacteria have evolved a number of diverse acclimation or regulatory strategies to withstand iron deficiency and optimize their photosynthetic performance. Under iron stress, cyanobacterial cells undergo chlorosis (loss of total Chl per cell), reduce the abundance of photosynthetic reaction centers and PBSs content (Öquist, 1974; Guikema and Sherman, 1983; Chen et al., 2018). On the other hand, in response to iron stress, various cyanobacteria species synthesize Chl-binding iron-stress-induced A (IsiA) proteins. IsiA is homologous to PsbC – the CP43 core antenna protein of PSII (Bibby et al., 2001; Boekema et al., 2001) and also known as CP43′. Apart from iron stress, it is also induced and required for growth under other stress conditions, such as high-light, salt, heat and oxidative stress (Vinnemeier et al., 1998; Singh et al., 2004; Havaux et al., 2005; Kojima et al., 2006).

IsiA is found in multiple copies that associate in rings around monomeric, trimeric or tetrameric PSI (Bibby et al., 2001; Boekema et al., 2001). The typical PSI–IsiA supercomplex consists of a closed ring of 18 IsiA around a PSI trimer (Bibby et al., 2001; Boekema et al., 2001). Under prolonged iron starvation, IsiA-PSI supercomplexes are formed with different numbers of IsiA and PSI monomers (Yeremenko et al., 2004; Kouřil et al., 2005), including larger ones where PSI is surrounded by double IsiA rings — with 18 IsiA monomers in the inner ring and 25 in the outer — forming a larger IsiA−PSI supercomplex (Chauhan et al., 2011). In addition to associating with PSI trimers, IsiA has been found to form rings and aggregates of its own (Ihalainen et al., 2005; van der Weij-de et al., 2007).

IsiA has been proposed to have a light-harvesting role - as an accessary antenna of PSI (Burnap et al., 1993) as well as a photoprotective one by dissipating excess energy. In iron-starved *Synechocystis* sp. PCC 6803, IsiA can increase the effective absorption cross section of PSI by 60% (Ryan-Keogh et al., 2012). On the other hand, the accumulation of IsiA aggregates that display much faster excitation decay than other membrane-bound Chl-protein complexes suggests that they serve as a thermal sink (Park et al., 1999; Ihalainen et al., 2005). The mechanism of quenching in IsiA aggregates is not yet determined (Chen et al., 2017). It has also been proposed that IsiA functions as a storage pool for Chls, holding up to 50% of the cellular Chl content during transition into iron limitation (Singh and Sherman, 2007; Schoffman and Keren, 2019). During recovery IsiA provides an accessible reservoir to support the rebuilding of the photosynthetic apparatus.

The structure of a PSI–IsiA complex from *Synechocystis sp.* PCC 6803 has been determined by cryoelectron microscopy (Toporik et al., 2019). The IsiA monomer resembles CP43 with six transmembrane helices but lacks the distinct loop connecting helices V and VI. It coordinates 17 Chls, 13 of which occupy similar positions to CP43 (Toporik et al., 2019), and four carotenoids — three at the IsiA–IsiA interface and one between IsiA and PSI. It coordinates Chls bridging IsiA subunits in the ring and also plays a role in the interaction between IsiA and PSI. Based on a detailed analysis of the X-ray structure of PSI, the PsaF subunit was assigned as a major recognition and interaction site for IsiA (Fromme et al., 2003). It was shown that lack of PsaF and PsaJ subunits results in the accumulation of smaller, partial IsiA rings (Kouřil et al., 2003). The recent higher-resolution structure confirms that, on the stromal side, the C-terminus of IsiA interacts with PsaF, PsaJ and PsaK (Toporik et al., 2019).

The capacity of PSI to accept energy from IsiA in the PSI–IsiA supercomplexes depends on the fast photochemical trapping of energy in the PSI core – 20–50 ps depending on species – and on the energetic coupling between Chls in the IsiA ring and the PSI core. Time-resolved spectroscopy studies of isolated PSI–IsiA complexes from *Synechocystis sp.* PCC 6803 and *Synechococcus sp.* PCC 7942 have shown that the addition of IsiA increases the trapping time from 20–25 ps to 39–44 ps (Melkozernov et al., 2003; Andrizhiyevskaya et al., 2004), which suggests fast and highly efficient energy transfer from IsiA to PSI. However, a wide range of energy transfer times is reported – from under 2 ps (Melkozernov et al., 2003) to 180 ps at 77 K (Akita et al., 2020). Moreover, these values are based on in vitro studies but information on the efficiency and dynamics of cyanobacterial energy transfer in vivo is scarce. The variety of PSI–IsiA and IsiA complexes present in thylakoid membranes makes it difficult to evaluate which of its purported roles – light-harvesting, Chl storage or energy dissipation - is more dominant. Finally, it is not clear how the efficiency of energy transfer depends on the oligomerization state of PSI and the presence or absence of the PsaF/PsaJ subunits.

In this work, we applied steady-state and time-resolved fluorescence spectroscopy to probe IsiA-PSI energy transfer in iron-starved *Synechocystis sp.* PCC 6803 cells with trimeric PSI (WT), monomeric PSI (Δ*psaL* mutant) and monomeric PSI lacking PsaF and PsaJ subunits (ΔFIJL mutant). We show that IsiA is capable of transferring energy in vivo to all types of PSI but to a different extent and that the absence of PsaL, and especially PsaF/J subunits leads to diminished IsiA levels in the cells under iron deficiency, thereby demonstrating IsiA’s primary role as an accessory light-harvesting antenna to PSI.

## MATERIALS AND METHODS

### Growth conditions and preparation

*Synechocystis* sp. PCC 6803 (hereafter referred to as *S.* 6803) cells were grown photoautotrophically in BG-11 medium under continuous white fluorescent light (∼35 μmol photons m^−2^ s^−1^) at 30°C. The wild type strain (culturable under light-activated heterotrophic growth and maintained in our lab), the Δ*psaL* mutant obtained in the same WT background (Kłodawska et al., 2015) and the subunit-depleted ΔFIJL mutant (Malavath et al., 2018) were used in this study. Iron-stressed cultures were obtained by inoculating the cells in BG-11 medium lacking iron-containing compounds. The cultures were grown for one week. Thylakoid membranes were prepared as described in Akhtar et al. (2021). PSI–IsiA and IsiA complexes were isolated according to Yeremenko et al. (2004) with small modifications. The thylakoid membranes were solubilized by incubating with 1 % *n*-dodecyl-β-maltoside (β-DDM) on ice. The unsolubilized material was removed by centrifugation at 30,000 g for 30 min. The supernatant was filtered through 0.45 µm filters and loaded on an ion-exchange chromatography column (Hi-Trap Capto Q ImpRes, Cytiva, USA). The fractions containing PSI–IsiA were eluted using a 0–300 mM Mg_2_SO_4_ gradient, concentrated, and further purified using a size-exclusion chromatography column (HiPrep 16/60 Sephacryl S-300 HR, Cytiva, USA).

### Chlorophyll content determination

Chls were extracted from the cell suspensions in 90% methanol and the Chl contents were determined spectrophotometrically using molar absorption coefficients described in Lichtenthaler (1987).

### Immunoblotting analysis

Suspension of thylakoid membranes containing 1 µg Chl was mixed with 6× Laemmli buffer (375 mM Tris/HCl [pH 6.8], 60% [v/v] glycerin, 12.6% [w/v] sodium dodecyl sulfate, 600 mM dithiothreitol, 0.09% [w/v] bromophenol blue) and incubated at 75°C for 10 min before loading. Proteins were fractionated on sodium dodecyl sulphate-polyacrylamide gel electrophoresis (SDS-PAGE) and immunoblotted with the corresponding polyclonal antibodies (produced in rabbits) purchased from Arisera AB: IsiA (AS 06 111), 1:1000 and PsaA (AS 06 172), 1:2000. The SuperSignal West Pico PLUS Chemiluminescent Substrate (Thermo Scientific) was used to detect these proteins with horseradish peroxidase-conjugated, anti-rabbit secondary antibody (Bio-Rad #1706515).

### Absorption and CD spectroscopy

Absorption and CD spectra in the range of 350–750 nm were recorded at room temperature with an Evolution 500 dual-beam spectrophotometer (Thermo Fisher Scientific, USA) and a J-815 spectropolarimeter (Jasco, Japan), respectively. The measurements were performed in a standard glass cuvette of 1-cm optical path length with 1 nm spectral bandwidth. The absorption spectra of the cells were corrected for the baseline by using bleached cells as a reference. CD spectra in UV region were measured at the B23 CD beamline of the Diamond synchrotron (UK). For synchrotron-radiation CD measurements, the samples were placed in 0.2 mm quartz cuvettes.

### Steady-state fluorescence spectroscopy

Fluorescence emission spectra in the visible range were measured at room temperature and 77K using a FP-8500 (Jasco, Japan) spectrofluorometer. The samples were diluted to absorbance of 0.1 per cm at the red maximum. Emission spectra in the range of 620–780 nm were recorded with excitation wavelength of 440 nm and 580 nm and excitation/emission bandwidth of 2.5 nm. The measurements were performed with 1 nm increment and 1s/ 4s integration time for room temperature and 77 K respectively. For measurements at 77 K, samples were cooled in an optical cryostat (Optistat DN, Oxford Instruments, UK) using liquid nitrogen. The spectra are corrected for the spectral sensitivity of the instrument using a calibrated light source as a reference (Jasco ESC-842).

### Time-resolved fluorescence spectroscopy

Picosecond time-resolved fluorescence measurements were performed with a time-correlated single-photon counting instrument (FluoTime 200/PicoHarp 300 spectrometer, PicoQuant, Germany) equipped with a microchannel plate detector (R3809, Hamamatsu, Japan). Excitation was provided by a Fianium WhiteLase Micro (NKT Photonics, UK) supercontinuum laser, generating white-light pulses with a repetition rate of 20 MHz. Excitation wavelength of 440 nm was used to excite selectively Chls. The fluorescence decays were recorded at wavelengths of 608–744 nm with 8 nm steps, at room temperature, and 605–750 nm with 5 nm steps at 77 K. All the samples were diluted to an absorbance of 0.03 at excitation wavelength. For the room temperature measurements, the suspension (whole cells or isolated thylakoids) was placed in 1 mm flow cell and circulated at a flow rate of 4 ml/min. For 77 K measurements, the suspension was placed in a 1 mm demountable cryogenic quartz cell and cooled in an optical cryostat (Optistat DN, Oxford Instruments, UK). The total instrument response (IRF) measured using 1% Ludox as scattering solution has width of 40 ps. The data are corrected for the spectral response of the detector. Global multiexponential lifetime analysis with IRF reconvolution was performed using MATLAB.

## RESULTS

### Fluorescence emission spectra of iron-starved cells

The main chlorophyll-protein complexes found in the cyanobacterial thylakoid membranes – PSI, PSII and IsiA show distinct fluorescence emission spectra at 77 K – IsiA has an emission maximum at 686, PSII at 686 nm and 696 nm and PSI around 724 nm (Andrizhiyevskaya et al., 2002). This difference can be used to estimate the relative abundance of IsiA from the 77 K emission spectra of iron-starved cells. A potential drawback of the method is that it is not possible to tell whether IsiA fluorescence changes are caused by alteration of its concentration in the cell or by changes in its fluorescence lifetime or energy transfer efficiency. Nevertheless, a semiquantitative comparison can be made.

The fluorescence emission spectra of control (+Fe) and iron-starved (−Fe) cells of WT and mutant *S.* 6803 cells are compared in Figure 1 (note that all spectra are normalized to the intensity of the PSI emission maximum at 724 nm). The fluorescence spectra of iron-starved cells show the expected increase of emission at 686 nm that corresponds to the accumulation of IsiA (Öquist, 1974). However, the 686 nm emission is markedly different in the iron-starved Δ*PsaL* and ΔFIJL mutants compared to the WT, whereas no such differences are present in control (+Fe) cells. Consistent with the fluorescence spectra, the absorption spectra of iron-stressed cells show an increased absorption at 678 nm that is more prominent in WT than in the two mutants (Supplementary Fig. S1). From these data, we can hypothesize that the accumulation of IsiA under conditions of iron deficiency depends on the oligomeric state of PSI and the presence of the PsaF subunit.

**Figure 1.**
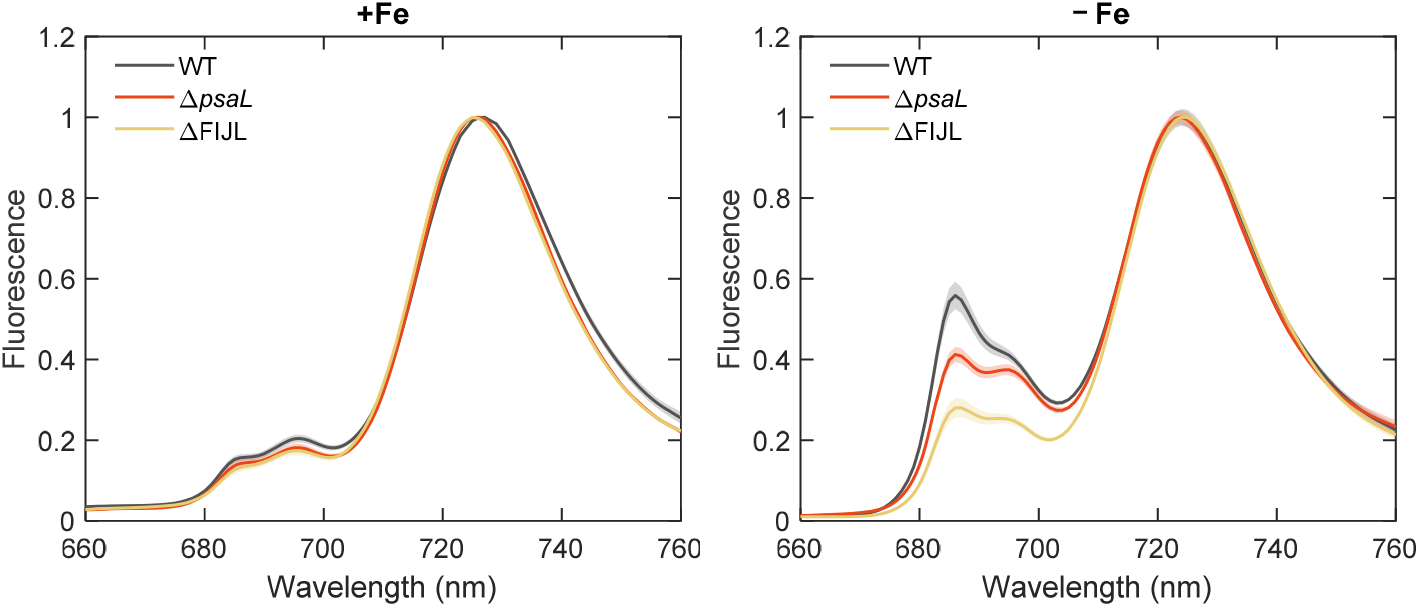
Fluorescence emission spectra of intact cells of WT *Synechocystis* sp. PCC 6803, Δ*psaL* and ΔFIJL mutants grown in regular BG-11 medium (+Fe) and in a medium devoid of iron (−Fe). The spectra are recorded at 77 K with 440 nm excitation and normalized to the maximum at 724 nm. The +Fe spectra are average of five and the −Fe spectra are average of 10–13 different batches with standard errors represented by the shaded area.

Based on the fluorescence intensities at 696 and 686 nm, we can estimate the PSII and IsiA emission relative to the PSI emission intensity at 724 nm (Table 1). All strains subjected to iron limitation show approximately two-fold increased PSII:PSI emission ratio compared to the control cells grown in iron-replete medium, which is consistent with a reduced PSI content in the cells. The emission of IsiA in both strains with monomeric PSI was significantly lower relative to the IsiA emission in WT cells – 49% and 37% for Δ*psaL* and ΔFIJL, respectively (with a standard error of 12%).

**Table 1.**
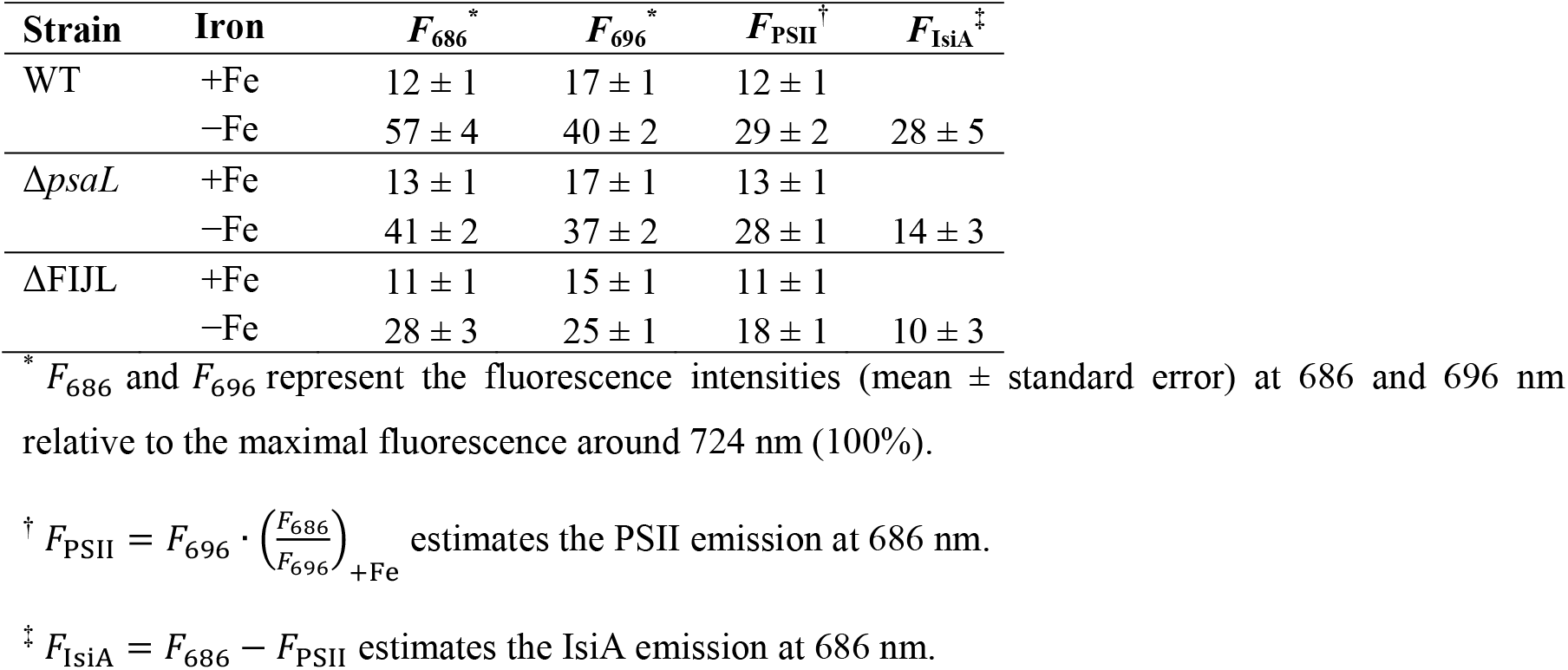
Peak analysis of the 77K emission spectra of *S.* 6803 cells

The fluorescence spectra recorded with 580 nm excitation allow us to compare the PBSs emission in the control and iron-starved cells (Supplementary Fig. S2). In control BG-11 medium, the spectra show that the monomeric mutants accumulate lower amounts of PC (645 nm peak) but have stronger PSII emission as compared to the WT, as reported previously (Akhtar et al., 2022). Iron-starved cells of both mutant strains show similar differences in the phycocyanin region. In contrast, both Δ*psaL* and ΔFIJL show lower emission in the PSII/IsiA region than WT. This result is also consistent with lower IsiA accumulation in the Δ*psaL* and ΔFIJL strains.

Because intact cells may be prone to artefacts from light scattering and reabsorption of the emitted fluorescence, we have performed the same experiments with isolated thylakoids from control and iron-starved cells. The fluorescence emission spectra were very similar to their counterparts from intact cells (Supplementary Fig. S3 and Table S1) and from the fluorescence intensities at 686 and 696, we estimated approximately the same relative IsiA emission in the mutant thylakoids (49% and 37% for Δ*psaL* and ΔFIJL relative to IsiA emission in WT thylakoids, standard error 10%).

These data strongly suggest that the IsiA content in iron-stressed *S.* 6803 cells depends on the oligomerization state of PSI and further on the presence of the PsaF/J subunits. However, it should be noted that the relative fluorescence intensities cannot be generally equated with the concentrations of IsiA in the cells.

### CD spectra of thylakoid membranes and PSI–IsiA complexes

As an additional probe for the changes in abundance of IsiA, we compared the CD spectra of isolated PSI–IsiA complexes and of thylakoid membranes isolated from iron-stressed cells (Figure 2 and supplementary Fig. S4). The thylakoid membranes were solubilized with detergent to suppress the light scattering. IsiA shows a negative-amplitude CD band at 438 nm. In contrast, thylakoids from cells grown in control BG-11 medium have near-zero CD signal at 438 nm and no local minimum in this wavelength range, which makes the 438 nm band a convenient indicator for IsiA.

**Figure 2.**
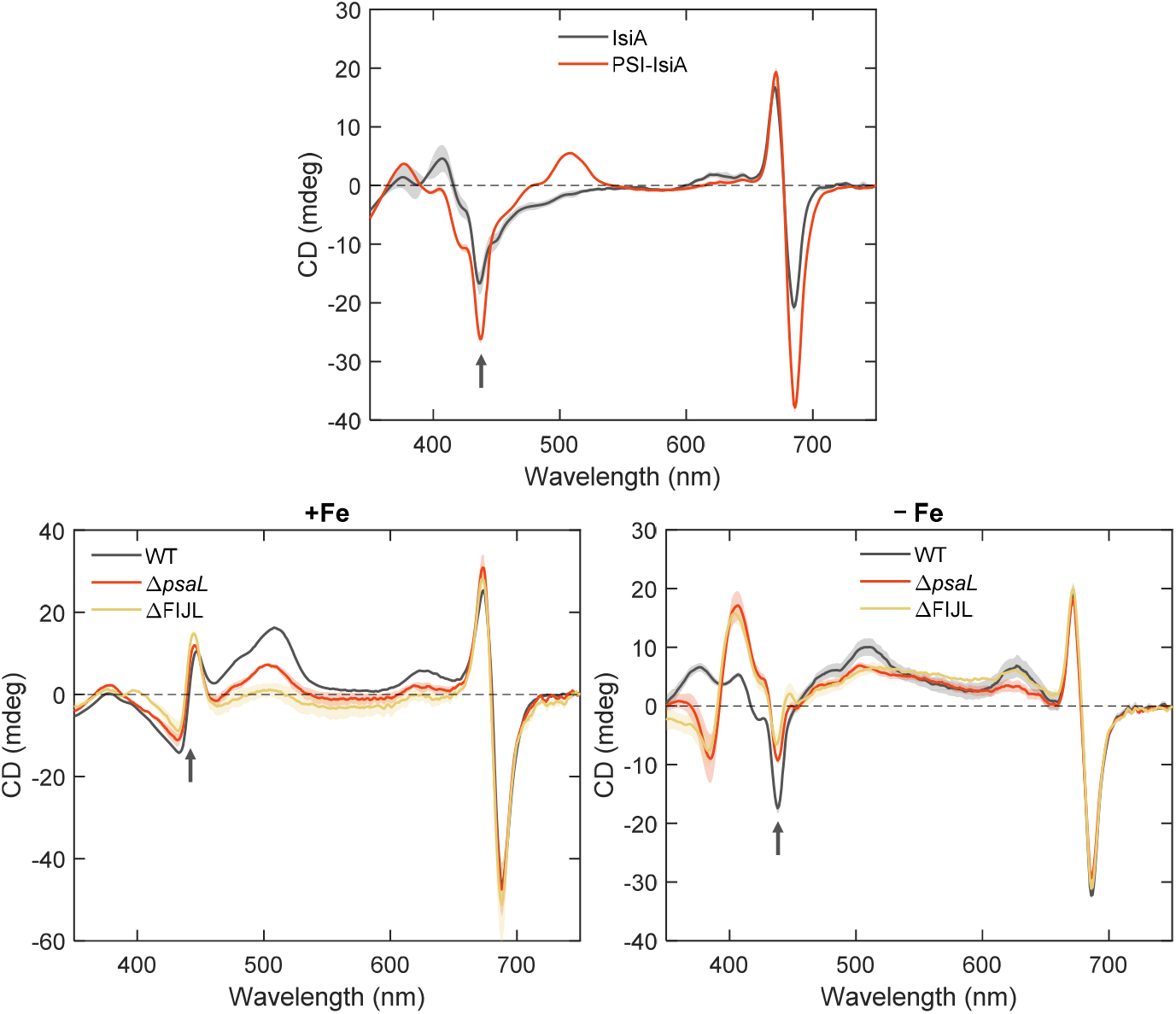
Room-temperature circular dichroism spectra of IsiA and PSI–IsiA complexes isolated from WT *S.* 6803 and of thylakoid membranes of control (+Fe) and iron-stressed (−Fe) *S.* 6803 WT, Δ*psaL* and ΔFIJL, measured in buffer containing 0.03% β-DM. The spectra are normalized to unity absorbance at the Chl Q_y_ maximum. The arrows indicate the IsiA-specific CD band at 438 nm. The figures show average spectra from 3–4 independent measurements on different batches with standard errors represented by the shaded area.

In line with the fluorescence emission data, we find that the IsiA CD peak is significantly diminished in the monomeric PSI mutants. The mutants also show CD changes in the UV region, which are of unknown origin. We take the differential CD amplitude at 448 nm and 438 nm as representative of IsiA (as it is negligible in control thylakoids). Considering that the CD signal is proportional to the respective chromophore concentration, it can be estimated that thylakoids of Δ*psaL* and ΔFIJL contain respectively 57 ± 8% and 48 ± 9% IsiA as compared to WT.

### Immunoblotting quantification

As a further test for the IsiA content of iron-starved *S.* 6803, we performed immunoblotting analysis with antibodies against IsiA and the PsaA subunit of PSI (Figure 3). As expected, in iron deficiency conditions IsiA accumulated in WT cells at the expense of photosystems (PsaA). Both the Δ*psaL* and ΔFIJL mutants showed lower relative IsiA content and higher PSI content compared to the WT. On a total Chl basis, the estimated IsiA content was 58% and 45% for Δ*psaL* and ΔFIJL compared to WT – in excellent agreement with the CD spectroscopy results.

**Figure 3.**
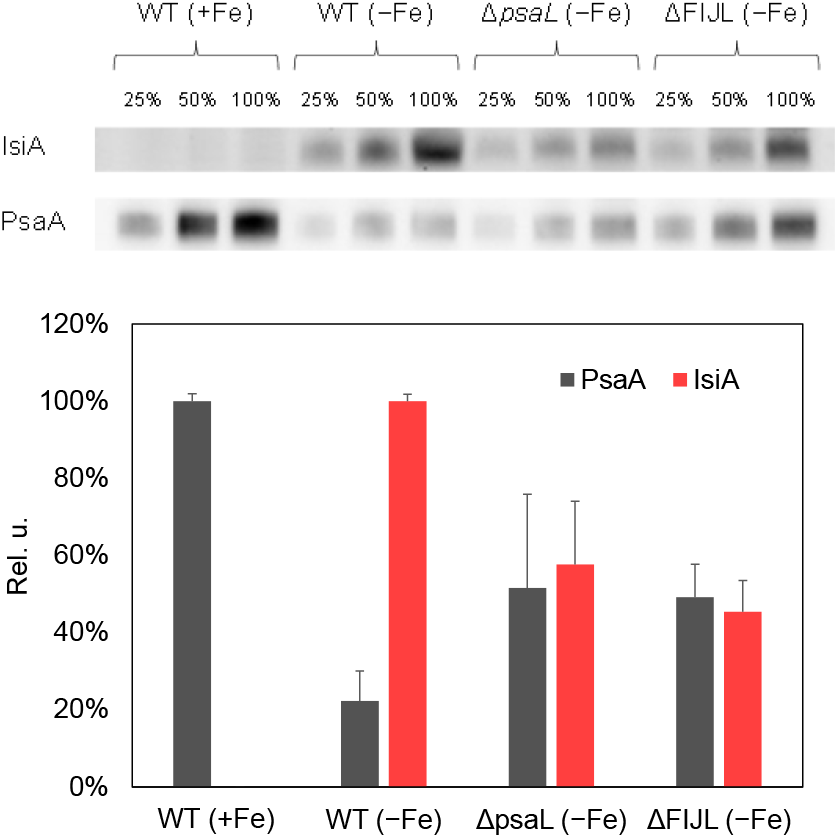
Immunobloting analysis of PSI and IsiA content. Top - representative immunoblots to monitor the changes in IsiA and PsaA levels in thylakoid membranes isolated from control *S.* 6803 WT cells (+Fe) and from WT, Δ*psaL* and ΔFIJL cell cultures grown in iron-deficient medium (−Fe). Each gel lane was loaded with 0.025, 0.05 or 0.1 μg Chl (100%). The faint background in the IsiA bands of control (+Fe) samples are probably due to reactivity of the IsiA antibodies with PSII (CP43). Bottom – averaged PsaA and IsiA concentrations (relative to WT) estimated from densitometric analysis of immunoblots. Standard errors from 3 replicates are indicated. Densitometric analysis was done using GelAnalyzer 19.1 (www.gelanalyzer.com) by Istvan Lazar Jr., PhD and Istvan Lazar Sr., PhD, CSc.

### Fluorescence kinetics of PSI–IsiA

The steady-state spectroscopy data show that the concentration of IsiA in iron-stressed *S.* 6803 is affected by PSI abundance. Next, we performed time-resolved fluorescence spectroscopy measurements to probe the efficiency of energy transfer in the intact cells. More specifically, we aimed to determine the dynamics of energy transfer from IsiA to PSI in the intact cells and whether IsiA can effectively function as a light-harvesting antenna of PSI in the absence of the PsaL and PsaF subunits.

The fluorescence kinetics were measured at room temperature in the range of 610–740 nm with 440 nm excitation wavelength, which excites predominantly Chl. Global multiexponential analysis of the fluorescence decays was applied to obtain fluorescence decay lifetimes and decay-associated emission spectra (DAES). The fluorescence kinetics of isolated IsiA and PSI–IsiA complexes can be described with three DAES (Figure 4). IsiA fluorescence decays with lifetimes of 70 ps, 0.5 ns and 2.5 ns. The shorter-lived component shows the well-known fluorescence quenching of IsiA aggregates (Ihalainen et al., 2005), whereas the longer-lived components may originate from monomeric IsiA, as evidenced by the blue-shifted DAES.

**Figure 4.**
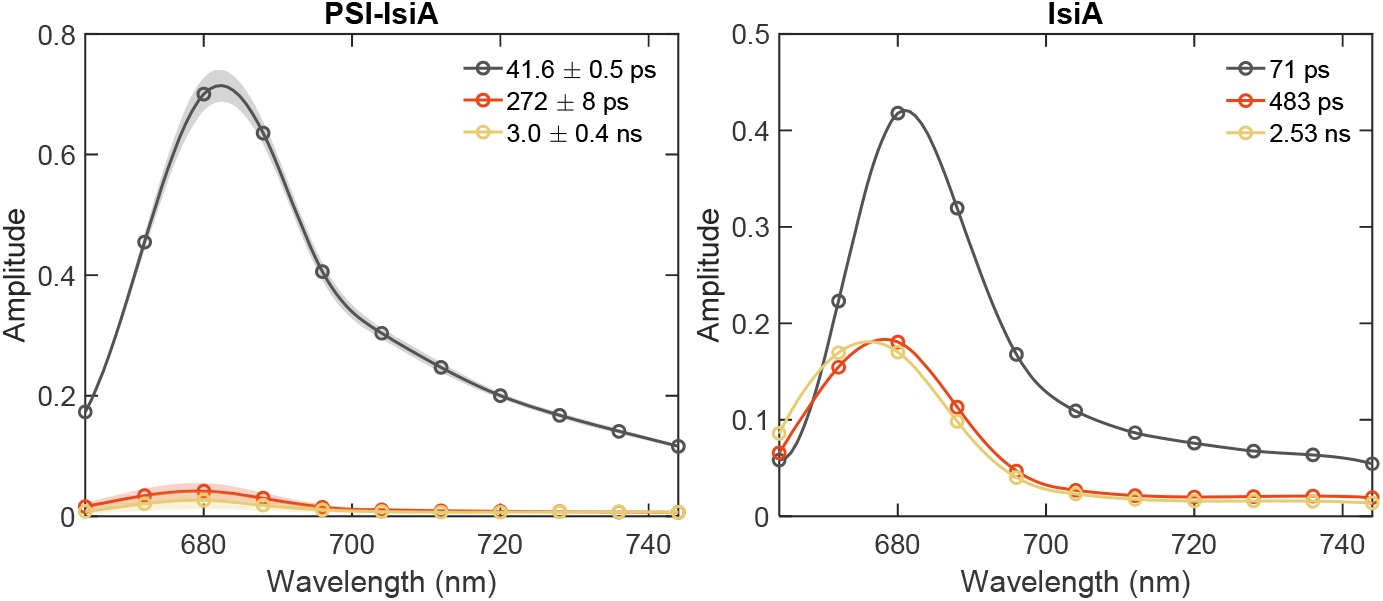
DAES of PSI–IsiA and IsiA complexes isolated from *S.* 6803 (WT) cells grown in iron-deficient medium. The spectra are obtained by global lifetime analysis of the fluorescence decays recorded at room temperature with 440 nm excitation. The spectra of PSI–IsiA are averages of 3 independent replicates. The shaded areas represent standard errors.

The fluorescence of PSI–IsiA complexes decays with a main lifetime of 42 ps. The two longer-lived components with lifetimes of 272 ps and 3 ns likely represent uncoupled IsiA complexes and Chls. The total trapping time of PSI–IsiA (42 ps) is significantly longer than that of isolated PSI alone (26 ps, see Supplementary Fig. S5), evidently due to energy migration between IsiA and PSI. We can roughly estimate the energy migration time τ_mig_ by considering the relationship τ_tot_ = τ_tr_ + τ_mig_, where τ_tr_ is the trapping time of PSI alone and τ_tot_ is the total decay time of PSI–IsiA. It follows that the IsiA-PSI energy transfer time in these complexes is approximately 16 ps, in good agreement with published results (Melkozernov et al., 2003; Andrizhiyevskaya et al., 2004; Chauhan et al., 2011).

Figure 5 compares the DAES for the fluorescence kinetics measured from control and iron-starved cells. The fluorescence kinetics of control cells (+Fe) can be described with three decay lifetimes, plus an additional ns component with a negligible amplitude. The fastest-decaying component with a lifetime of ∼25 ps belongs to PSI, whereas the two slower-decaying components with lifetimes of 170 ps and 550 ps (in WT) show the decay of PSII and partially PBS fluorescence. Detailed description of the kinetics can be found elsewhere (Akhtar et al., 2021). In comparison with the control, the amplitudes of the two long-lived components increase three-fold in iron-stressed cells, which can be attributed to emission from IsiA complexes. Moreover, the short-lived decay components (35–42 ps) have lifetimes and DAES that are similar to that of PSI–IsiA supercomplexes. Similar results in terms of fluorescence lifetimes and DAES features were obtained from thylakoid membranes isolated from control and iron-starved cells (Supplementary Fig. S6).

**Figure 5.**
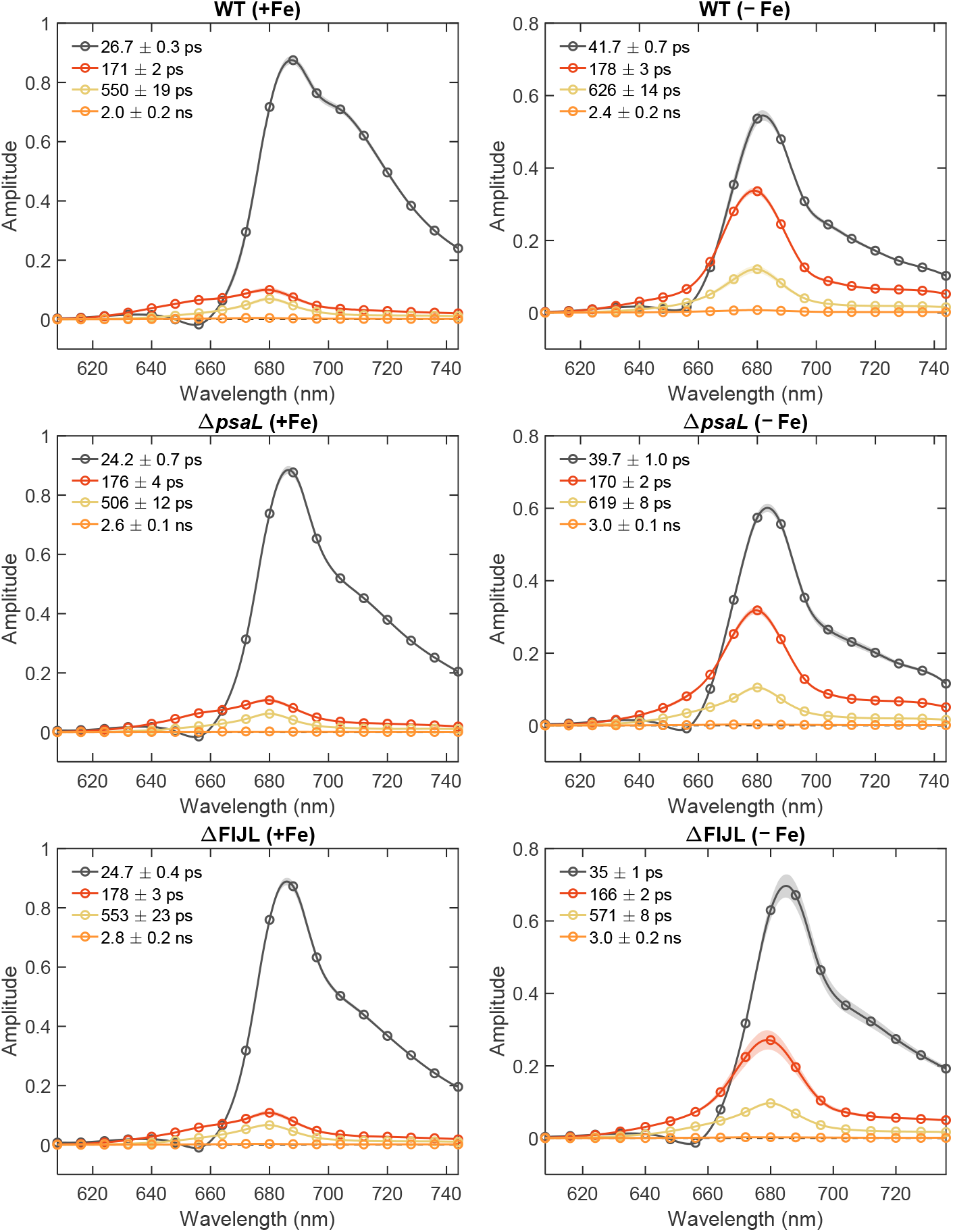
DAES of intact *Synechocystis* cells (WT, Δ*psaL* and ΔFIJL mutants) grown in regular BG-11 medium (+Fe) and in a medium devoid of iron (−Fe), obtained by global lifetime analysis of the fluorescence decays recorded at room temperature with 440 nm excitation. The spectra are averages of 7–11 independent replicates. The shaded areas represent standard error.

We ascribe the 35–42 ps DAES to PSI–IsiA supercomplexes (cf. Figure 4) and the longer-lived decay components to energetically decoupled IsiA (and partially to PSII emission). Decay lifetimes in the 70–100 ps range are not resolved in intact cells, which may suggest that isolated IsiA aggregates are more strongly quenched than IsiA in the cells.

From the above we conclude that the iron-stressed cells of *S.* 6803 contain both PSI–IsiA supercomplexes and unconnected IsiA aggregates. The same applies to the two mutants. It is noticeable that the relative amplitudes of the long-lived DAES associated with free IsiA are lower in the mutants, in line with the overall lower emission and concentration of IsiA in these cells. For instance, the wavelength-integrated amplitude of the 170 ps IsiA component relative to the 40 ps PSI component is reduced by 30% in ΔFIJL compared to the WT (Supplementary Table 2). The PSI–IsiA lifetime is also slightly shorter in the mutants, especially in ΔFIJL – 35 ps compared to 42 ps in WT. From the difference of the PSI lifetimes in control cells and the PSI–IsiA lifetimes in iron-starved cells, we estimate that the migration time in WT cells is about 15 ps – almost equal to that of the isolated PSI–IsiA complexes from the same strain, whereas for Δ*psaL* and ΔFIJL it is 10 ps. The shorter apparent migration time may indicate that PSI monomers that lack the PsaF subunit connect fewer IsiA units than PSI monomers with the PsaF subunit present.

### Quantification of IsiA from time-resolved fluorescence

To estimate the fractions of IsiA complexes connected to PSI and of free IsiA aggregates, we apply kinetic model fitting to the DAES. The average number of IsiA in the PSI–IsiA complexes determines the effective trapping lifetime and the shape of the PSI–IsiA DAES – more IsiA complexes result in stronger emission from IsiA and a longer trapping lifetime and vice versa. Unconnected IsiA complexes mainly contribute to the amplitude of the longer-lifetime components (170 ps and 600 ps). Following this, the sizes of both IsiA pools can be found by a simple fit, as illustrated in Figure 6**Error! Reference source not found.** and Supplementary Fig. S7 (more details are available in the Supplementary Information).

**Figure 6.**
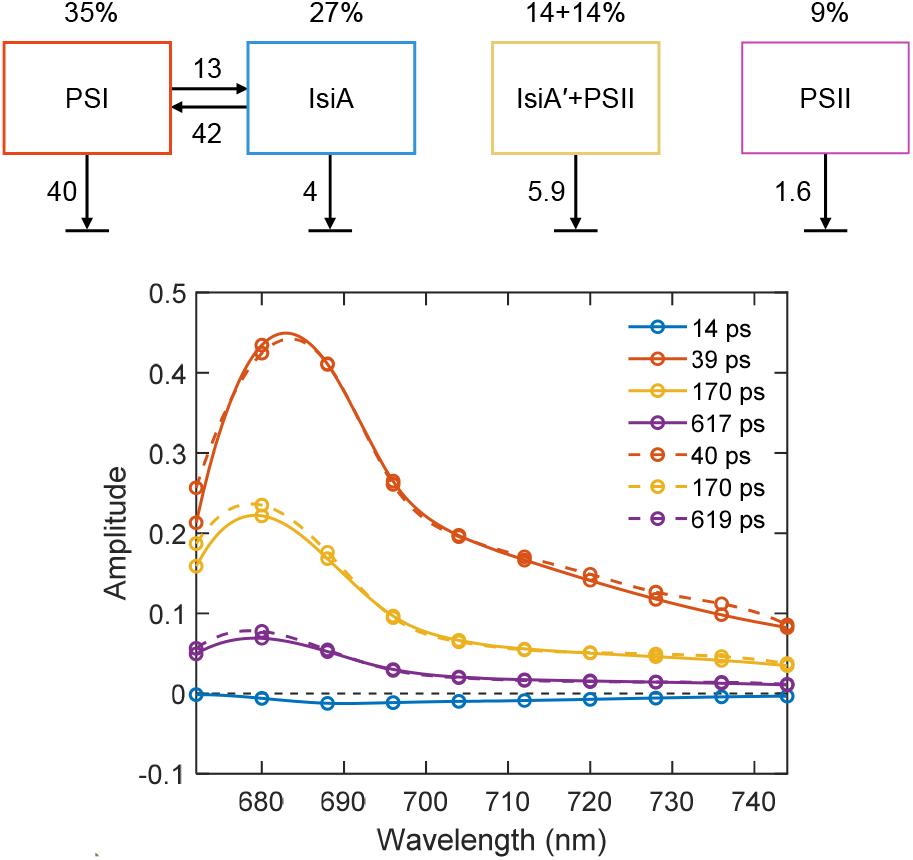
Kinetic model of PSI–IsiA fluorescence in *Synechocystis* Δ*psaL* cells. The four compartments represent PSI–IsiA complexes, unconnected IsiA and PSII. The decay and energy transfer rate constants (ns^−1^) are indicated next to the arrows. The numbers above the compartments indicate the relative excitation of each compartment, proportional to the number of Chls. The PSI and IsiA emission spectra are modelled using the area-normalized DAES obtained from isolated complexes and PSII is represented by the PSII-associated DAES (500 ps) of iron-replete cells. The graphs show the model DAES (solid lines), compared with the DAES obtained from global lifetime analysis of the measured fluorescence kinetics of Δ*psala* cells.

The results are summarized in Table 2 for the three *S.* 6803 strains. The analysis reveals that the relative PSI content (per Chl basis) of the monomeric strains is higher than the WT, whereas the relative IsiA content is lower – in agreement with the immunoblotting analysis (Figure 3). Furthermore, we estimate 6–7 IsiA complexes connected to PSI in WT cells, which is consistent with 18 IsiA per PSI trimer (Kouřil et al., 2003). Note that the model was validated on isolated PSI–IsiA complexes, for which the fitting yielded an IsiA:PSI molar ratio of 6:1, consistent with the known structure (Toporik et al., 2019). In WT cells, we find additional 4–5 unconnected IsiA per PSI monomer to a total IsiA:PSI molar ratio of about 11:1. This value is in close agreement with the results of Fraser et al (2013) who estimated a molar IsiA:PSI ratio of 12 in *S.* 6803 grown under similar iron-limited conditions. In the monomeric PSI strains, we find that both the connected and unconnected IsiA pools are reduced compared to WT cells. The total IsiA:PSI molar ratios were 58% and 38% for Δ*psaL* and ΔFIJL cells, respectively. The PSI subunit knockouts appeared to affect both the connected and the free IsiA pools to a similar extent.

**Table 2.**
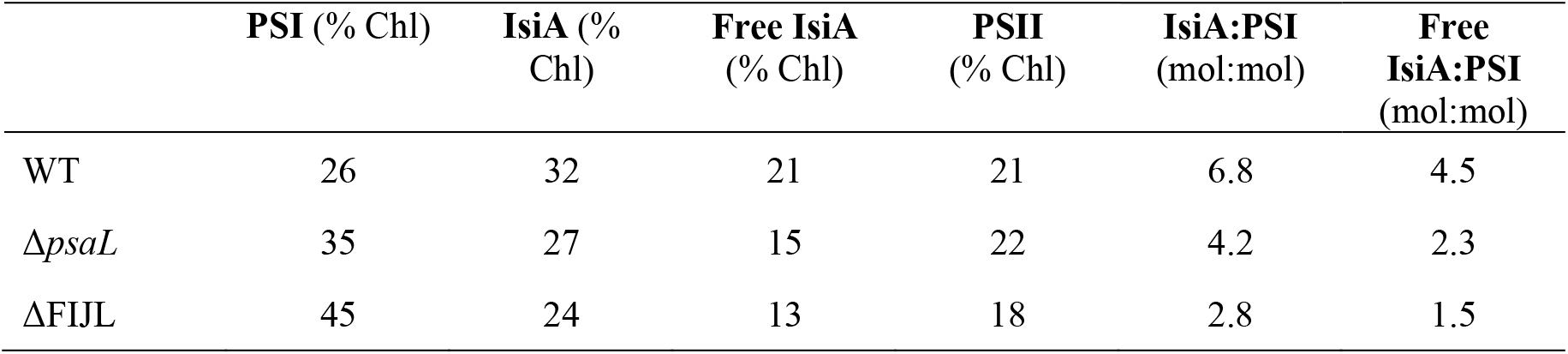
Quantification of IsiA based on time-resolved fluorescence

|  | PSI (% Chl) | IsiA (% Chl) | Free IsiA (% Chl) | PSII (% Chl) | IsiA:PSI (mol:mol) | Free IsiA:PSI (mol:mol) |
| --- | --- | --- | --- | --- | --- | --- |
| WT | 26 | 32 | 21 | 21 | 6.8 | 4.5 |
| $\Delta psaL$ | 35 | 27 | 15 | 22 | 4.2 | 2.3 |
| $\Delta FIJL$ | 45 | 24 | 13 | 18 | 2.8 | 1.5 |

### Low-temperature fluorescence kinetics

The DAES at 77 K (Figure 7) show that IsiA is connected to PSI and that excitation energy transfer can be described with two lifetimes – around 30 ps and 80–90 ps. The ΔFIJL mutant has lower amplitudes of these decay components at 680 nm, consistent with fewer IsiAs connected to PSI. This result alone cannot explain the large decrease in the 680 nm fluorescence intensity in the steady-state emission spectra. In addition, we observe non-transferring IsiA that decays with 300 ps. This component is significantly diminished in the mutant compared to the WT. Therefore, the lower fluorescence intensity in the steady-state spectra of the iron-stressed mutants compared to the WT is due the reduced number of IsiA complexes, both connected to PSI and, to a larger extent, the highly fluorescent non-transferring IsiA.

**Figure 7.**
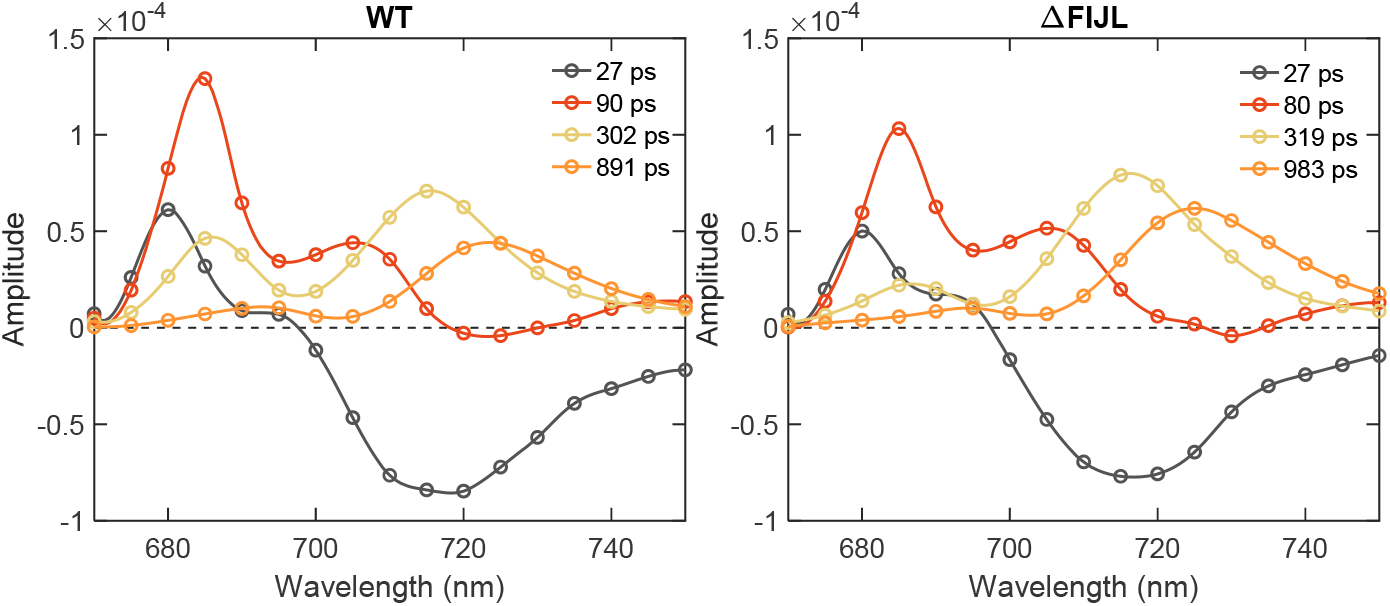
Representative DAES of *Synechocystis* WT and ΔFIJL mutants cells grown in a medium devoid of iron (−Fe), obtained by global lifetime analysis of the fluorescence decays recorded at 77 K temperature with 440 nm excitation.

## DISCUSSION

### Trimeric PSI forms functional supercomplexes with IsiA in vivo

The capability of many species of cyanobacteria to form rings of IsiA (CP43′) under iron stress has been known for more than two decades (Bibby et al., 2001; Boekema et al., 2001). It is also well established that the IsiA subunits in isolated PSI–IsiA supercomplexes are functionally connected to the PSI core via efficient energy transfer (Melkozernov et al., 2003; Andrizhiyevskaya et al., 2004; Chauhan et al., 2011). There is no reason to doubt that a similar energetic connectivity also occurs in vivo. The efficient energy transfer allows IsiA to increase the absorption cross-section of PSI, allowing it to trap more photon energy per unit time, in this way reducing the number of Fe-rich PSI complexes necessary to maintain photosynthesis. Nevertheless, several authors have suggested that IsiA has functions other than to facilitate PSI light harvesting – such as dissipating excess light energy, maintaining PBSs and storing Chls until more favourable conditions (see Chen et al., 2018 and refs. therein).

This study investigates the physiological role of IsiA by evaluating its ability to transfer energy to PSI in intact cells of iron-starved *S.* 6803. To this end we compare the picosecond Chl fluorescence kinetics of cells and of isolated PSI–IsiA complexes, as a reference. The isolated complexes showed that excitations are trapped on a timescale of 42 ps. Similar trapping times were found in PSI–IsiA from *S.* 6803 by (Andrizhiyevskaya et al., 2004). From the difference between the trapping time in the supercomplex and the PSI core, it follows that the effective timescale of energy equilibration between IsiA and the PSI core is about 15–16 ps, which is close to the 10 ps lifetime of energy transfer determined by transient absorption spectroscopy (Melkozernov et al., 2003). We found that the PSI kinetics in intact WT *S.* 6803 cells subjected to iron deficiency was remarkably similar to that of isolated PSI–IsiA. Not only was the main trapping lifetime identical (42 ps) but the corresponding DAES had virtually identical shape, that can be approximated as the sum of the PSI core (60%) and IsiA (40%) emission. The similarity strongly suggests that the kinetics of energy transfer in PSI–IsiA in vivo is the same as in isolated state and that the isolated PSI–IsiA sample is representative of the naturally occurring structural organization. Time-resolved fluorescence spectroscopy of cells at 77 K revealed two lifetimes of IsiA–PSI transfer in cells – 20–30 ps and 80–90 ps. Similar lifetimes were observed in isolated PSI–IsiA at 77 K (Akita et al., 2020). These result further supports the functional equivalence of PSI–IsiA complexes isolated in vitro and in the intact thylakoid membrane.

The structural robustness of the PSI–IsiA supercomplex observed in vitro (Chauhan et al., 2011) is the basis for resilient energy transfer. From these results it can be assumed that the primary physiological role of IsiA is light harvesting. However, the time-resolved fluorescence data (at 77 K and room temperature) suggest that a sizeable population of IsiA (approximately 40% of the total IsiA) are not functionally coupled to PSI. These results support electron microscopy studies on iron-stressed cyanobacteria that have found the presence of IsiA aggregates unconnected to PSI in cells (Yeremenko et al., 2004). The iron-stressed cells have significantly increased emission at 680 nm on equal Chl basis, similar to the results of Schrader et al. (2011) with *S.* 6803 grown in iron-deficient nutrient-rich media. The increase in fluorescence was attributed to the existence of IsiA aggregates that are not connected to reaction centres.

### Dual role of IsiA

In line with previous studies, we show that under iron stress, cells can accumulate IsiA in excess of what is needed for functional light harvesting by PSI and it serves dual function. It has been seen by electron microscopy that IsiA builds supercomplexes without PSI under prolonged iron stress (Yeremenko et al., 2004). We observe both IsiA connected to PSI and unbound IsiA aggregates in a ratio of approximately 1.5:1 in WT *S.* 6803. According to in vitro studies, unconnected IsiA aggregates provide protection for PSII from photooxidation (Yeremenko et al., 2004; Havaux et al., 2005; Ihalainen et al., 2005; van der Weij-de et al., 2007). Similar to LHCII in higher plants, which has a dual light-harvesting/photoprotective role and forms supercomplexes of different sizes with and without PSII, IsiA is suggested to play a dual role, increasing the absorption cross-section of PSI on the onset of iron stress and regulating and balancing the light harvesting to protect PSII from overexcitation by shading (Yeremenko et al., 2004; van der Weij-de et al., 2007). The significant size of the unconnected IsiA pool (∼40% of the total IsiA) appears to support such a role.

### The accumulation of IsiA depends on the PSI architecture

The functional significance of the trimeric organization of PSI in cyanobacteria has been puzzling. Recently it was shown that the oligomeric state has an effect on the abundance and architecture of PBS and the distribution of excitation energy harvested by PBSs between PSII and PSI (Akhtar et al., 2022). The two mutant strains of *S.* 6803 with monomeric PSI – Δ*psaL* and ΔFIJL had lower phycocyanin content and their PBS delivered less excitation energy to PSI, indicating that the trimeric PSI organization facilitates PBS–PSI energy transfer. In this study we subjected those mutant strains of *S.* 6804 to iron-limiting conditions to investigate how the PSI architecture and subunit composition affects the accumulation of IsiA and the IsiA–PSI energy transfer.

All experimental approaches – steady-state and time-resolved fluorescence, CD spectroscopy and immunoblotting show that the monomeric PSI mutants have significantly reduced IsiA content compared to the WT. All analyses show that the IsiA content decreases in the order WT > Δ*psaL* > ΔFIJL. Therefore, the accumulation of IsiA in iron-stressed cells depends not only on the presence of the PsaL subunit but also on the PsaF/PsaJ subunits. Kouřil et al. (2003) showed that the PSI–IsiA supercomplexes are smaller in the absence of PsaF/PsaJ subunits and proposed that these subunits are needed for the binding of IsiA to PSI. In agreement with the microscopy studies, we find that PSI in the ΔFIJL mutant binds less than half the number of IsiA complexes than WT PSI. We conclude that the accumulation of IsiA in the thylakoid membrane is controlled by its ability to functionally connect to the PSI trimers; however, the exact feedback mechanism that regulates the expression of IsiA remains to be elucidated.

It could be expected that if IsiA does not form stable PSI–IsiA supercomplexes with the mutant PSI, a larger fraction will remain unconnected to PSI in the mutant cells. This, however, does not seem to be the case, as the mutants have equally reduced pools of connected and unconnected IsiA compared to the WT *S.* 6803 strain and approximately equal ratios of connected/free IsiA (Table 2). We can speculate that in the strains with subunit-depleted PSI, IsiA cannot sufficiently enhance the PSI absorption cross-section and, consequently, the cells maintain higher relative PSI content. More experiments are needed to test whether the altered acclimation strategy in the mutants will significantly hamper their capacity to grow under prolonged iron limitation conditions.

### Conclusions

To summarize, this study shows that IsiA formed functional supercomplexes in iron-stressed cells where energy was efficiently transferred to PSI in a manner similar to isolated PSI–IsiA complexes. Subunit-depleted PSI was unable to accommodate the same number of IsiA as the trimeric WT PSI, resulting in a smaller effective cross-section and leading to an overall lower IsiA content and higher PSI content on Chl basis. These results strongly support the light-harvesting role of IsiA in vivo as well as the role of the PsaF/PsaJ subunits in mediating PSI–IsiA interaction. At the same time, the observation of significant fractions of IsiA that are not functionally connected to PSI in all three *S.* 6803 strains suggests that light harvesting is not the only functional role of the complex under iron limitation conditions.

Since both mutants with monomeric PSI displayed significantly reduced ability to functionally bind IsiA, this study represents further piece of evidence for a specific physiological role of the oligomerization of PSI in cyanobacteria, in addition to the recently demonstrated role of trimeric PSI in facilitating energy transfer from PBS (Akhtar et al., 2022).

The findings shed light on IsiA’s dual function in light harvesting and photoprotection in cyanobacteria under iron stress conditions. The results also demonstrate the possibility for quantitative spectroscopic detection of IsiA and its functional state in live cyanobacterial cells, that could be utilized to develop specific biosensors for assessing iron bioavailability and iron limitation stress in the field (Schrader et al., 2011). Further research can explore the specific mechanisms and regulatory factors involved in the formation of PSI–IsiA supercomplexes and the dynamics of IsiA aggregation, contributing to a deeper understanding of the complex interplay between light harvesting and photoprotection in photosynthetic organisms.

## Acknowledgements

We thank Prof. Dario Leister for the gift of the ΔFIJL mutant of *S.* 6803.

The work was supported by grants from the National Research, Development and Innovation Fund (NKFI FK-139067 to PA and 2018-1.2.1-NKP-2018-00009 to PHL) and the Еötvös Loránd Research Network (SA-76/2021 to PA). CD measurements at the B23 beamline of the Diamond Light Source Ltd. were supported by the project CALIPSOplus under Grant Agreement 730872 from the EU Framework Programme for Research and Innovation HORIZON 2020.

## Author contributions

PA and PHL conceptualized the study; PA performed most experiments. FBG maintained cyanobacterial cultures, isolated thylakoid membranes, and participated in spectroscopic measurements. SK and SZT designed and performed the immunoblotting study. All authors have contributed to the writing of the manuscript and given approval to the final version.

